# Foraging at night under artificial light: consequences on reproductive senescence and lifetime reproductive success for a diurnal insect

**DOI:** 10.1101/2023.03.02.530765

**Authors:** Gomes Elisa, Lemaître Jean-François, Rodriguez-Rada Valentina, Débias François, Desouhant Emmanuel, Amat Isabelle

## Abstract

1. The increasing use of artificial light at night (ALAN) is currently a major anthropogenic disturbance, with largely unappreciated eco-evolutionary consequences for nocturnal but also diurnal organisms. It has been hypothesized that light pollution could create new opportunities for the latter to forage and reproduce at night, which is called the ‘night-light’ niche, with fitness consequences still scarcely explored.
2. We exposed diurnal parasitoid wasps (*Venturia canescens*) to one of three light-at-night conditions: control (0 lux), low intensity (0.7 lux) or high intensity (20 lux) throughout their lives. We then monitored changes in both behavioural and life-history traits, namely daytime and nighttime feeding and egg-laying activity, reproductive senescence, lifespan and lifetime reproductive success.
3. Light pollution influenced the nighttime activity of wasps. The proportion of wasps feeding and laying eggs at night increased, and we also detected a tendency for a higher nighttime reproductive success under a high intensity of light pollution. Surprisingly, high intensity of light pollution also increased the wasps’ lifespan. Such changes did not affect the lifetime reproductive success of the wasps, but influenced the distribution of ovipositions between day and night.
4. Reproductive senescence occurs in *V. canescens*, evidenced by the linear decline in daily reproductive success with age regardless of the light condition. ALAN conditions, in interaction with mother age, affected developmental time in offspring, highlighting an effect on reproductive senescence.
5. We demonstrated that light pollution induced the use of the ‘night-light’ niche in a diurnal insect, with a shift in the distribution of egg-laying events between day and night. While we did not observe strong consequences on individual fitness, such changes in the dynamics of parasitism behaviour may nevertheless have major consequences for population dynamics, especially in natural conditions.

## INTRODUCTION

Among human-induced environmental changes such as habitat loss, chemical pollution or climate change (Sih *et al*., 2011), artificial light at night (ALAN) is a rapid and pervasive phenomenon (Falchi *et al*., 2016). The rise of artificial light at night during the 19^th^ century and its strong increase with the development of electric lighting profoundly altered natural light cycles. Daily and seasonal cycles of light and dark, as well as lunar light cycles, are predictable environmental variations that act as major cues for organisms (Gaston *et al*., 2017; Kronfeld-Schor *et al*., 2013). Their disruption by artificial light at night therefore has multiple ecological and biological impacts for the living world (Sanders *et al*., 2020) as well as potentially important evolutionary consequences (Davies & Smyth, 2018; Desouhant *et al*., 2019; Swaddle *et al*., 2015).

Initially, studies on the detrimental effects of artificial light at night were solely focused on nocturnal organisms, such as bats, moths or fireflies, which were more likely to suffer from behavioural disturbance (Deora *et al*., 2021; Firebaugh & Haynes, 2016; Russo *et al*., 2017). However, diurnal organisms are also impacted by illuminated nights since artificial lighting creates new opportunities for diurnal species to forage or display reproductive behaviours at night (‘night-light’ niche, Longcore & Rich, 2004). So far, evidence of the use of the ‘night-light niche’ come mostly from diurnal vertebrate species, such as birds with an advanced onset of activity (Dominoni *et al*., 2014; de Jong *et al*., 2016) or an increase in nighttime activity (Ouyang *et al*., 2017). In the common swift (*Apus apus*), high intensity of light at night during the breeding season led birds to be active throughout the night instead of joining their nest (Amichai & Kronfeld-Schor, 2019). In the European blackbird (*Turdus merula*), individuals exposed to rather dim levels of ALAN (about 0.15 lux) extended their foraging activity in the evening compared to individuals in darker areas (Russ *et al*., 2015). Observations also showed that diurnal species can take advantage of light sources, which attract arthropods, to forage during the night (*e*.*g*., several bird species, Lebbin *et al*., 2007; two lizard species, *Anolis leachii* and *Anolis wattsi*, Maurer *et al*., 2019; an arthropod, *Platycryptus undatus*, Frank, 2009). Interestingly, diurnal arthropods can also use the night-light niche. For instance, a mesocosm study suggested that some diurnal parasitoid species may extend their parasitism activity into the night when exposed to low levels of ALAN (Sanders *et al*., 2018). In a subsequent paper on the same biological system, (Kehoe *et al*., 2020) showed that global parasitism rate increases with daylength and that ALAN intensifies this effect. Quantifying accurately the impact of artificial lighting on fitness-related traits at night and during the day is now essential to fully understand the ecological and evolutionary consequences of ALAN on diurnal organisms, and ultimately their interaction with other species.

Lifetime reproductive success (‘LRS’), defined as the number of offspring produced during an individual’s lifespan, is a keystone variable to estimate fitness of organisms (Clutton-Brock, 1988). Lifetime reproductive success depends on longevity and reproductive success, two traits that we examined in this study. In the last decade, empirical studies have highlighted that the decrease in reproductive success with age (*i*.*e*., reproductive senescence or reproductive ageing) is a pervasive process that occurs in both laboratory and wild populations (Nussey *et al*., 2013; Zajitschek *et al*., 2020). Reproductive senescence patterns can differ within species, notably in response to environmental conditions (Cooper & Kruuk, 2018; Tidière *et al*., 2016), which can be altered by human activities. Anthropogenic disturbances may therefore ultimately impact senescence patterns. Indeed, some studies suggested a potential link between artificial light at night and age-related changes in biological traits, potentially mediated by increased oxidative stress following the presence of light during the night (studies on laboratory rats, El-Bakry *et al*., 2018; Vinogradova *et al*., 2009) or disruption of melatonin cycles (Reiter *et al*., 2017). However, a direct influence of ALAN on reproductive senescence and LRS is still rarely studied (but see McLay *et al*., 2018).

The aim of this study was hence to test whether artificial light at night affected the pattern of reproductive senescence, longevity and lifetime reproductive success of a diurnal parasitoid wasp, *Venturia canescens*. We designed an experiment that allowed us to consider both nocturnal and diurnal foraging behaviours separately. Indeed, while *V. canescens* is a diurnal species showing no activity in the dark (in absence of any stimulus related to feeding and oviposition), it has previously been demonstrated that females exposed to ALAN moved during the night (Gomes *et al*., 2021). Artificial light at night could therefore allow the wasps to exploit the night-light niche (*i*.*e*., to forage at night), which could consequently boost their lifetime reproductive success. We thus predicted that wasps exposed to ALAN would feed and lay eggs at night, unlike wasps not exposed to light at night. In addition, ALAN has previously been shown to also alter the diurnal behaviours of females: wasps exposed to light at night preferred to search for hosts rather than for food during the day, and tended to produce more offspring when given a single occasion to parasitize a host aggregate (Gomes *et al*., 2021). We therefore predicted that wasps exposed to ALAN would have higher parasitism rate and thus a greater reproductive success during the day and at night (but with fewer offspring produced at night than during the day) compared to wasps without light at night. In addition, wasps exposed to ALAN seem to allocate more to immediate reproduction (Gomes *et al*., 2021), so we also predicted stronger reproductive senescence in these individuals (characterized by an earlier onset of senescence and/or a steeper decline in reproductive performance), as expected under resource-based allocation trade-offs (Lemaître *et al*., 2015).

## METHODS

### Biological model and rearing conditions

*Venturia canescens* (Gravenhorst) is a solitary endoparasitoid hymenopteran (Ichneumonidae). It attacks lepidopteran larvae, mainly Pyralidae (Salt, 1975), in their second to fifth instar (Harvey & Thompson, 1995). Parasitoid females use a mandibular gland secretion released by host larvae when feeding to locate them (Castelo *et al*., 2003). *V. canescens* is a pro-synovigenic species (Jervis *et al*., 2001), which means that the females emerge with an initial load of mature eggs but additional eggs are also produced throughout their adult life. The eggs are hydropic (*i*.*e*., yolk-deficient) which means that they do not store nutrients (Le Ralec, 1995). Eggs are therefore not energetically costly to produce (Pelosse *et al*., 2011) and the lifetime reproductive success of *V. canescens* is expected to mostly depend on their longevity (approximatively 3 weeks in the lab) and ability to find and parasitize hosts. Feeding influences longevity (wasps feeding every 48h lived six times longer than wasps feeding at longer intervals, Desouhant *et al*., 2005), it should therefore increase the fecundity of *V. canescens* by giving females more opportunities to parasitize hosts (Harvey *et al*., 2001).

We used wasps of a thelytokous (*i*.*e*., parthenogenesis in which females produce only daughters from unfertilized eggs) strain established from about 70 wild females trapped near Valence (southern France) during the summer of 2016. The strain was maintained under controlled conditions (25±1°C, 60±5% relative humidity, 12:12 light:dark) in boxes containing larvae of *Ephestia kuehniella* (Lepidoptera: Pyralidae) as hosts and organic wheat semolina as feeding medium for the host larvae. The adult wasps were fed *ad libitum* with honey diluted 1:1 with distilled water, because in natural conditions *V. canescens* has frequently access to nectar or honeydew as food sources (Desouhant, Lucchetta, *et al*., 2010).

### Light conditions

We have created three experimental groups of wasps differing in their intensity of exposure to light at night. The wasps were kept in three thermostated chambers (SANYO Electric Co., Ltd, model number: MLR-351H) providing a temperature of 25°C and a relative humidity of 60%. Light cycle (12:12 h light:dark) and daytime light intensity (3500 lux) provided by neon tubes were similar in all three chambers. Each chamber was equipped with white LED lights (ribbon of LED SMD 5050, 6000-6500K, Sysled) on the ceiling to provide either a high intensity of light at night (20 lux, equivalent to the intensity of a street lamp; hereafter ‘high ALAN’ condition) or a low intensity of light at night (0.7 lux, equivalent to the intensity of a city skyglow, Bennie *et al*., 2016; hereafter ‘low ALAN’ condition). These lighting systems were switched on in two of the three chambers, the third being used for the ‘control’ condition (0 lux, total darkness at night). We measured light intensities to the nearest 0.01 lux with an illuminance meter (T-10MA, Konica Minolta^®^) before the experiment, and checked light intensity every time we switched light conditions between thermostated chambers (see below). Light measurements in the chambers were performed inside the clear plastic boxes containing the insects and subsequently used in the experiments. We alternated the light conditions between chambers every two weeks to prevent any “chamber effect”.

### Experimental set-up

We aimed to quantify the daily number of offspring produced by the wasps (exploiting host patches during the day and at night), as well as their feeding behaviour. We therefore designed an experimental set-up that made it possible to control when the wasps could feed and lay eggs, day and night. Host and food patches (see below) were placed on a Plexiglas^®^ plate in a standardised way (‘bottom plate’ in Figure S1 (A) in the Supplementary data). Right above, the wasps were individually kept in clear plastic boxes on a white Plexiglas^®^ plate (‘superior plate’ in Figure S1 (B) in the Supplementary Data), which was drilled with holes whose size and position coincide with the location of the host and food patches on the bottom plate. That superior plate could be lowered to allow the wasps to access the patches through the holes or lifted to prevent them to do so (Figure S2 (B) and (A) in the Supplementary Data). Lifting the superior plate also enabled to replace host patches with new ones when needed (see ‘Description of patch replacement’ in Supplementary Data). A coarse mesh was stretched under the superior plate to prevent the wasps from escaping through the holes when the superior plate was lifted (see Figure S2 in the Supplementary Data). That mesh had no effect on the wasp ability to lay eggs or acquire food (pers. obs. on 10 wasps, data not shown).

#### Preparation of host and food patches

Host patches consisted of 15 third-instar larvae in a Petri^®^ dish (55 mm diameter) filled with semolina and covered by a piece of organza. They were prepared from 7 to 10 days before they were used to allow the semolina to become impregnated with host kairomones.

Food patches were prepared by soaking cotton wools with 8 ml of sucrose solution (40%) in a Petri^®^ dish (35 mm diameter). The sucrose solution was dyed blue with food dyed (1% v/v), so we could easily determine if the insects ate or not by looking at the colour of their abdomen (see also in Desouhant *et al*., 2005). Food patches were prepared daily, just before their use in the experiment.

### Effect of ALAN on nighttime feeding behaviour

Under the three nighttime light conditions, the wasps had access to food either during the day or at night, alternately, followed by a 24-hour period without food (see Table 1). This experimental design aimed to test whether the wasps fed during daytime or nighttime when exposed to light. The 24h-period without food was planned to prevent a state of satiety that would have prevented the wasps from feeding on the following days.

**Table 1.** Schedule of the periods when the wasps were allowed to feed during a two-week experimental session. “∅” meant that food was not available. This sequence of feeding (D0-D3) was repeated until day 13.

| Day | Daytime<br>(8 a.m. to 8 p.m.) | Nighttime<br>(8:20 p.m. to 8 a.m.) |
| --- | --- | --- |
| Day 0 | Food | Ø |
| Day 1 | Ø | Food |
| Day 2 | Ø | Ø |
| Day 3 | Food | Ø |

### Effect of ALAN on lifetime reproductive success and reproductive senescence

Our experimental design consisted of three two-week experimental sessions, during which we recorded longevity and reproductive success of 76 wasps exposed to the three nighttime lighting conditions (*N* = 26, 26 and 24 in the control, low ALAN and high ALAN conditions, respectively) to estimate reproductive senescence patterns as well as lifetime reproductive success.

For every experimental session, we took newly emerged wasps, and individually provided them with a new host patch every day (from 8 a.m. to 8 p.m.) and every night (8:20 p.m. to 8 a.m. the next day) until death. There was a dark habituation period from 8 p.m. to 8:20 p.m., to prevent any behaviour due to the handling of the experimental set-up. Host larvae were concealed in the substrate (here, wheat semolina) and each host patch used during the nighttime period was exposed to ALAN for 12 hours. It was therefore highly unlikely that artificial light at night had consequences on hosts in our experiment. Age of death was recorded for each wasp, and those who were not yet dead after two weeks were integrated as censored individuals in the survival analysis (*N* = 12 out of 76 wasps in total; there was 3, 2 and 7 censored wasps in the control, low ALAN and high ALAN conditions, respectively). Wasps were then kept at -20°C to measure their body size, because this trait can positively influence longevity and reproductive success in *V. canescens* (Pelosse *et al*., 2011). We measured the tibia length as a proxy of body size as commonly done in this species (*e*.*g*., Harvey & Vet, 1997). For each wasp, we took a picture of the left hind tibia under a binocular magnifier coupled with a camera and measured its length to the nearest 0.01 mm with the software Motic Image Plus 2.0 (Motic, Hong Kong). We also measured the egg load at death by counting the mature eggs under a microscope. There was no difference in egg load at death between light conditions (data not shown). This means that potential differences in offspring production between light conditions were not due to egg limitation and variation in egg maturation during our experiments. We therefore used only body size as a covariate in analyses, to control for differences between individuals when assessing the effect of artificial light at night on lifetime reproductive success, longevity and reproductive senescence.

Each potentially parasitized host patch was kept in the rearing room (12:12 h light:dark, without light at night) and offspring emergences were counted for seven weeks (see the section ‘Dynamics of offspring production’ for a more detailed description). Classically, the development from egg to adult lasts 21 days in *V. canescens*. However, we extended the monitoring period because mother’s nighttime light condition, in interaction with mother’s age, has been shown to prolong offspring development time by up to 60 days (Gomes *et al*., 2021). Overall, our experimental design allowed to measure longevity and daily reproductive success of each wasp, distinguishing between daytime and nighttime egg-laying.

### Dynamics of offspring production throughout life

All host patches were monitored twice a week to count the total number of offspring per wasp and the dynamics of offspring production throughout the wasps’ lifespan. In addition, we also wanted to determine whether the dynamics of offspring emergence changed over the wasps’ lifespan, and whether these dynamics depended on the nighttime lighting conditions to which wasps had been exposed. For this purpose, we monitored the development time of offspring emerging from host patches parasitized by one-four-, seven- and ten-day-old wasps. We monitored the patches parasitized during both daytime and nighttime. Offspring emerging from these patches were frozen dry at -20°C to measure their body size (see protocol in the paragraph above).

### Statistical analysis

We performed the statistical analyses with R 4.0.2 (R Core Team, 2020). Throughout the statistical analyses, we used the packages *lme4* (Bates *et al*., 2014) and *nlme* (Pinheiro *et al*., 2019) to build linear models (‘LMs’), generalized linear models (‘GLMs’) and linear mixed models (‘LMMs’).

#### Effect of ALAN on lifespan and reproductive senescence

We investigated the effect of light condition on lifespan using survival analysis (package *survival*, Therneau, 2015). We fitted Cox proportional hazards models including light condition (factor with three modalities, ‘Control’, ‘Low ALAN’ and ‘High ALAN’), body size and thermostated chamber (factor with three modalities, ‘Chamber1’, ‘Chamber2’ and ‘Chamber3’), as well as their two-by-two interactions as explanatory variables. We selected the best model based on the Akaike Information Criterion (AIC). When models had a ΔAIC (*i*.*e*., difference between their AIC and the AIC of the best model) lower than 2, we selected the simplest model to satisfy parsimony rules (Burnham & Anderson, 2002). The full list of fitted Cox models is provided in supplementary data (Table S1). We used the package *rms* (Harrell, 2021) to compute post hoc comparisons between factor modalities in the best Cox model selected.

We tested the effect of light condition on reproductive senescence using three reproductive traits: the daily reproductive success (*i*.*e*., number of offspring produced from daytime and nighttime host patches), offspring quality (using body size as a proxy) and offspring development time (log-transformed to meet the assumption of normality). Age-related changes in these traits were assessed by building linear mixed models with the daily number of offspring, offspring body size or offspring development time as response variable, and mother’s age and light condition, as well as their interaction, as explanatory variables. For the daily number of offspring, we compared three types of age function: linear, quadratic and threshold. For the threshold function, the threshold value was determined by fitting a piecewise regression over a range of values between 1 and 14 days of age and selecting the value that gives the lowest residual deviance (Ulm & Cox, 1989). The best model selected had a threshold value set at 5 days. For offspring quality and development time, individuals whose mothers were 7 or 10 days old were grouped into the same age class of ‘[7-10] days’ because of small sample sizes, leading to three age classes. Age was therefore included as a factor (with three modalities, ‘1 day’, ‘4 days’ and ‘[7-10] days’) in the models for these two response variables. Mother’s identity was included in all the models as a random effect to account for confounding effects of pseudo-replication (sensu Hurlbert, 1984). Mother’s longevity was also included as a fixed effect to control for selective disappearance (*e*.*g*., if ‘low-quality’ individuals displaying a low reproductive success die younger, causing older age classes to display a high reproductive success) (Nussey *et al*., 2008). Finally, the total number of offspring produced was included as a covariate in the models for offspring body size and development time, to account for the potential trade-off between offspring quantity and quality (Berrigan, 1991; Godfray *et al*., 1991). The number of offspring for each mother’s age and light condition is given in Table 2, as well as the number of females by which these offspring were produced.

**Table 2.** Number of offspring in each cross-modality of artificial light at night (ALAN) condition and mother’s age. The number of mothers is written in brackets in the table.

|  |  | **Mother's age** |  |  |
| --- | --- | --- | --- | --- |
|  |  | *1 day* | *4 days* | *[7 – 10] days* |
| **Light condition** | *Control* | 121 (22) | 17 (5) | 12 (4) |
|  | *Low ALAN* | 98 (26) | 12 (9) | 10 (2) |
|  | *High ALAN* | 100 (21) | 31 (12) | 9 (5) |

As for Cox models, we selected the best model for daily reproductive success based on AIC. The full list of fitted models for daily reproductive success is provided in supplementary data (Table S2). For offspring quality and development time, the statistical significance of explanatory variables was based on the *F*-statistic with adjusted degrees of freedom (following Kenward & Roger, 1997).

For the analysis of nighttime behaviours and LRS, we built generalized linear models in which, unless otherwise specified, we included body size, light condition and thermostated chamber, as well as their two-by-two interactions, as explanatory variables. From the full models, we used a backward model selection approach to select the best model. When needed, we computed post hoc comparisons between factor modalities using the package *emmeans* (Lenth, 2020). Cohen’s *h* (Cohen, 1992) was computed to estimate effect sizes between proportions, using the *pwr* package (Champely, 2020). Magnitude of effect sizes was assessed according to the following thresholds: |*h*| < 0.2: negligible; 0.2 ≤ |*h*| < 0.5: small; 0.5 ≤ |*h*| < 0.8: medium; |*h*| ≥ 0.8: large (Cohen, 1992).

#### Nighttime feeding and egg-laying behaviours

To assess whether light condition influenced the wasps’ nighttime feeding behaviour, we fitted binomial GLMs (logit link). We computed the proportion of nighttime feeding events weighted by the total number of nights the individual had access to food. This response variable therefore took into account the wasp’s longevity. With regard to the egg-laying behaviour, we analysed the proportion of wasp that exploited host patches at night depending on light condition by fitting a binomial GLM with the binary variable ‘Patch exploitation behaviour at night’ (‘Yes’ or ‘No’) as response variable.

#### Effect of ALAN on lifetime reproductive success (LRS)

We analysed nighttime reproductive success (*i*.*e*., the total number of offspring that emerged from all the host patches exploited during the night throughout a wasp’s life) and daytime reproductive success (*i*.*e*., the total number of offspring that emerged from all the host patches exploited during the day throughout a wasp’s life) by fitting negative binomial GLMs (log link) that account for overdispersion in the data. LRS, estimated by pooling the number of offspring produced from daytime and nighttime host patches, was also analysed with a negative binomial GLM (log link). Finally, we analysed the distribution of egg-laying events between day and night by fitting a quasibinomial GLM (logit link).

## RESULTS

### Effect of ALAN on lifespan and reproductive senescence

#### Lifespan

While low and high ALAN conditions did not influence the lifespan in *V. canescens* compared to the control condition (Table 3, Figure 1), contrast analysis showed that the difference in lifespan between the low and high ALAN conditions was significant (*z*-value = 2.53, *P* = 0.01). The median longevity was 4.3 (95% CI = [2.9 ; 7.4]), 2.9 (95% CI = [2.9 ; 4.3]) and 6.1 (95% CI = [4.3 ; NA]) days in the control, low ALAN and high ALAN conditions, respectively. Body size also had a significant effect on the wasp longevity, with a larger body size reducing the risk of death by a factor of 0.008 (Table 3).

**Table 3.** Semi-parametric Cox proportional hazards (PH) model selected for the longevity data. The best model was selected based on Akaike Information Criterion (AIC). The full list of fitted models is provided in supplementary data (Table S1).

| Type | Variable | Coefficient ± SE<br>(hazard ratio) | z-value | P |
| --- | --- | --- | --- | --- |
| Semi-parametric Cox PH | Body size | -4.89 ± 1.3<br>(0.008) | -3.8 | <0.01 |
| | Light condition<br>(Low ALAN) | 0.47 ± 0.3<br>(1.60) | 1.5 | 0.13 |
| | Light condition<br>(High ALAN) | -0.35 ± 0.3<br>(0.70) | -1.1 | 0.3 |

**Figure 1.**
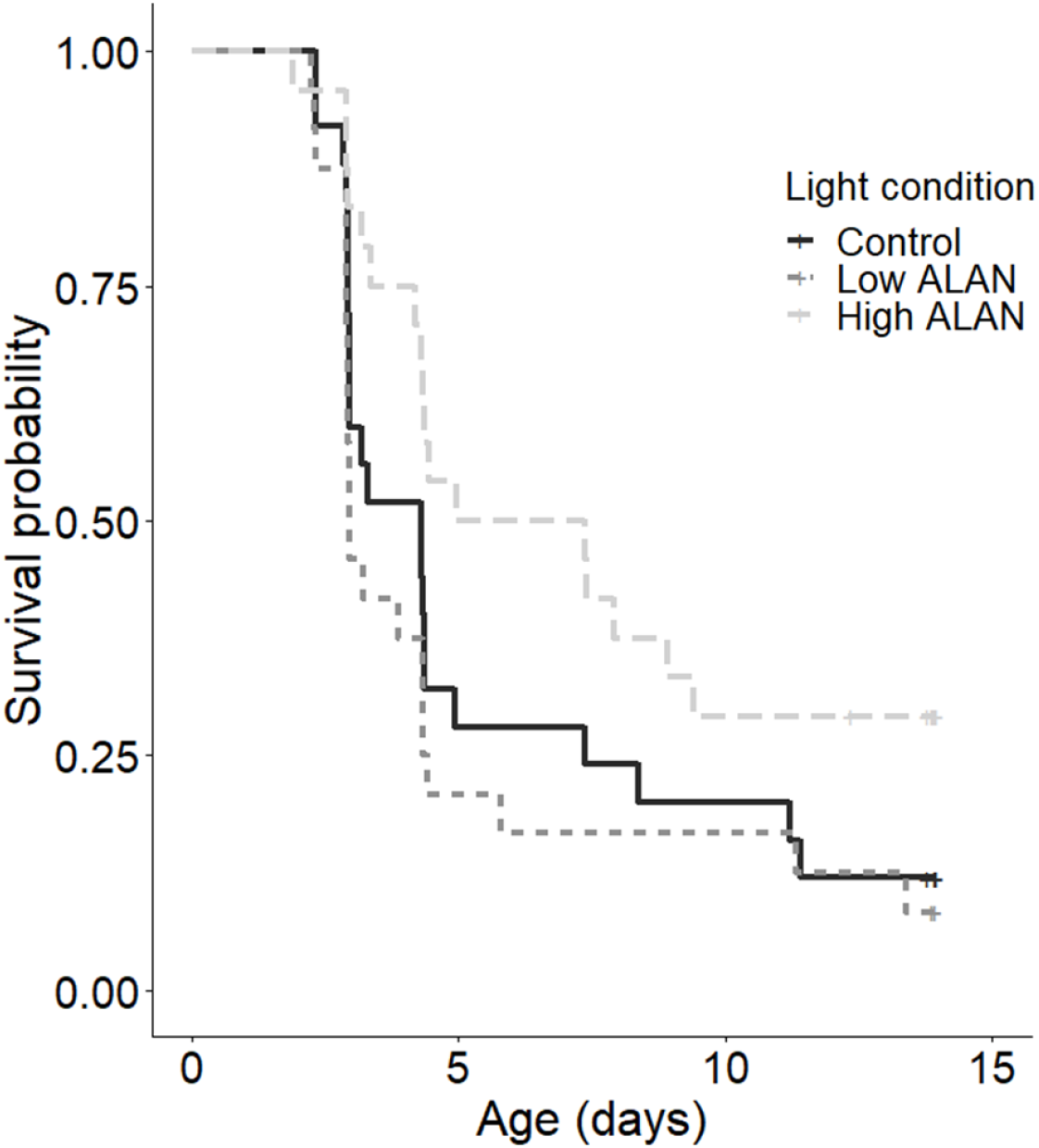
Survival curves for the control, low artificial light at night (ALAN) and high ALAN conditions.

#### Reproductive senescence

The best model selected to describe the changes of daily reproductive success with age included only a linear effect of age and no significant effect of light at night (Table 4 and Figure 2a).

**Table 4.** Linear mixed effects model selected for investigating senescence pattern in daily reproductive success. The best model was selected based on Akaike Information Criterion (AIC). The full list of fitted models in provided in supplementary data (Table S2).

| **Trait** | **Age function** | **Variable** | **Parameter estimate ± SE** | ***t*-value** |
| --- | --- | --- | --- | --- |
| Daily number of offspring | Linear | Age | -0.38 ± 0.04 | -9.74 |

**Figure 2.**
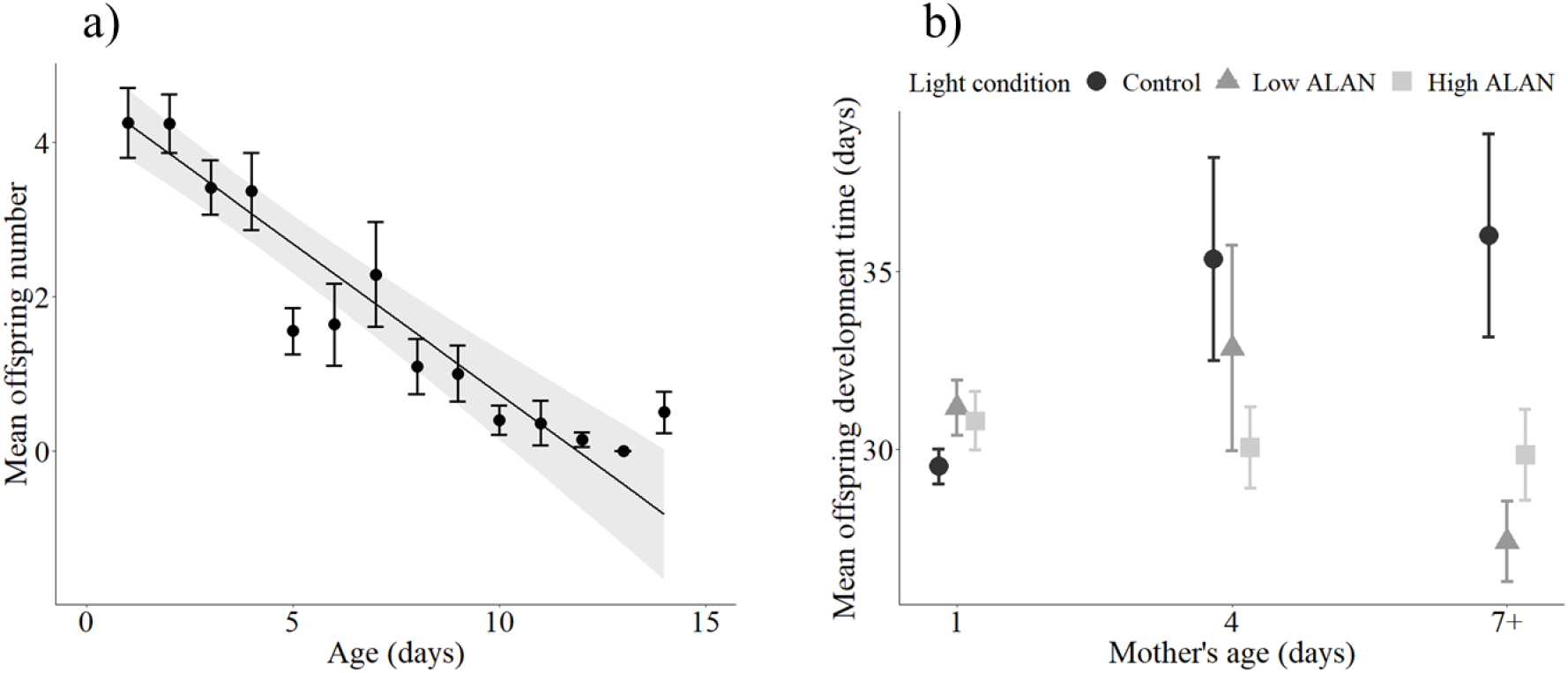
a) Age-related changes in reproductive traits in *V. canescens*. The points represent the average number of offspring produced per age and the bars correspond to Standard Error. The line represents prediction from the best selected model (linear effect of age) and the shaded area the 95% CI. b) Effect of mother’s age on offspring development time in the control, low ALAN and high ALAN conditions. Individuals older than 7 days of age were grouped into the ‘7+’ age class. Symbols represent mean ± SE.

Offspring size was not influenced by either mother’s age (*F* = 1.70, *df* = 2, 353.2, *P* = 0.18) or light condition (*F* = 0.68, *df* = 2, 63.6, *P* = 0.51) (Figure S3 in supplementary data). On the contrary, offspring development time depended on the interaction between mother’s age and light condition (*F* = 4.68, *df* = 4, 336.3, *P* < 0.01). More precisely, for offspring whose mothers were in the ‘7+’ age group, individuals in the control condition took significantly longer to develop than those in the low ALAN condition (about 9 days longer in average; post hoc test: *t-ratio* = 3.5, *P* < 0.01) (Figure 2b). However, the difference in development time was not significant between individuals in the control and high ALAN conditions (about 6 days in average; post hoc test: *t-ratio* = 1.8, *P* = 0.18), nor between individuals in the low and high ALAN conditions (about 2 days in average; post hoc test: *t-ratio* = -1.8, *P* = 0.18).

### Nighttime feeding and egg-laying behaviours

Light conditions influenced the proportion of nighttime feeding events (weighted by the number of nighttime feeding opportunities): 63%, 51% and 81% in the control, low and high ALAN conditions, respectively (*χ*² = 8.81, *df* = 2, *P* = 0.01; Figure 3). Post-hoc tests showed that the only significant difference was between low and high ALAN conditions (*z* = 2.80, *P* = 0.02) with a biological difference considered as medium (effect size: Cohen’s *h* = 0.65) (Figure 3b). However, the effect size between control and high ALAN conditions (Cohen’s *h* = 0.43) suggested that the difference, albeit non-statistically significant, was also moderately biologically relevant.

**Figure 3.**
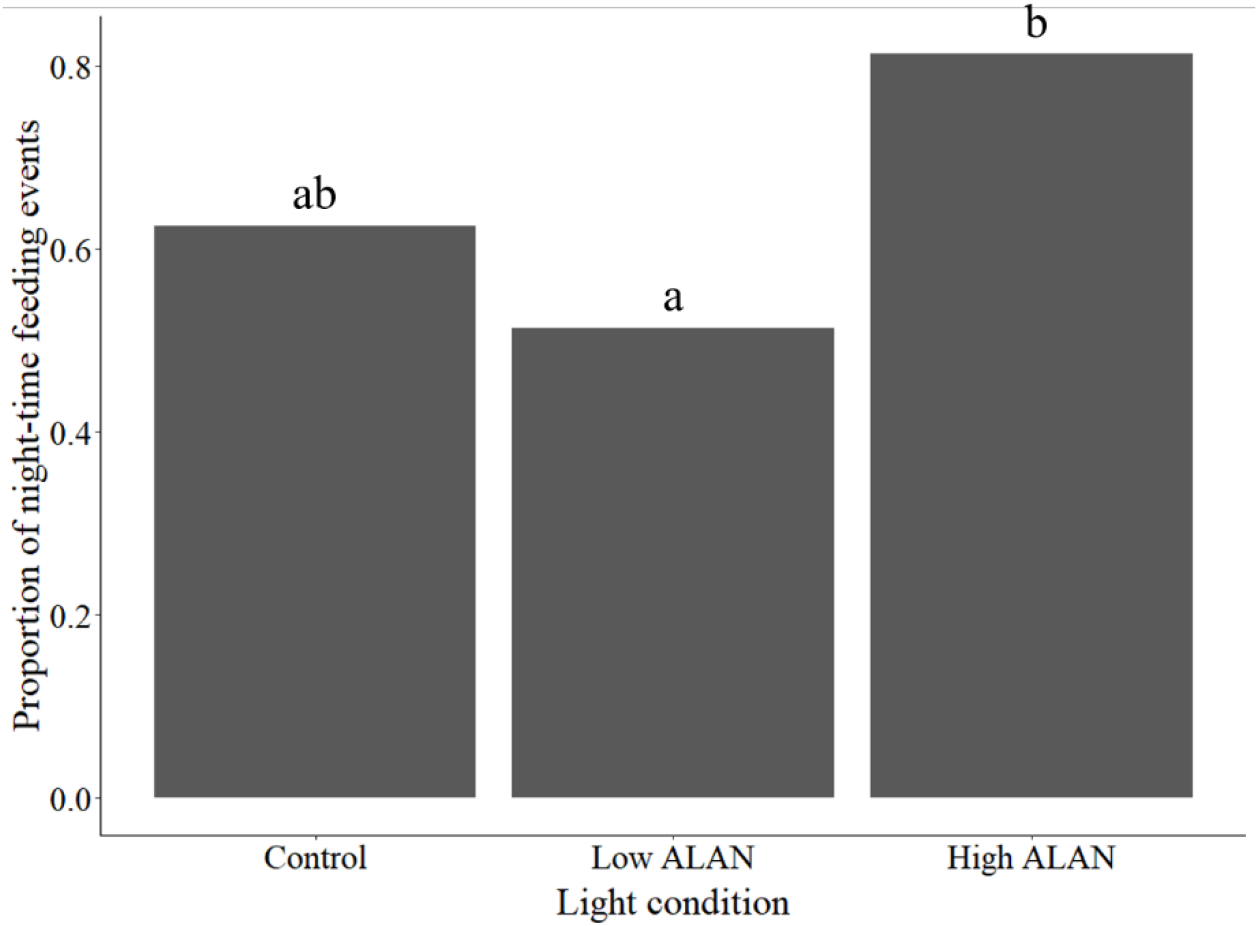
Proportion of nighttime feeding events (weighted by the number of nighttime feeding opportunities) for the control and artificial light at night (ALAN) conditions. Different letters show a significant statistical difference at the 0.05 level.

With regard to the nighttime egg-laying behaviour, light condition influenced the percentage of wasps that exploited host patches (*i*.*e*., that have had offspring) at night: 75%, 80% and 100% in the control, low and high ALAN conditions, respectively (*χ²* = 10.0, *df* = 2, *P* = 0.007). Effect sizes suggested a strong effect of high ALAN condition compared to control (Cohen’s *h* = 1) and low ALAN (Cohen’s *h* = 0.93) conditions.

### Effect of ALAN on lifetime reproductive success

Daytime reproductive success was not statistically different between light conditions (*χ*² = 2.17, *df* = 2, *P* = 0.34) (Figure 4). However, there was a tendency for light condition to influence the nighttime reproductive success (*χ* ² = 5.46, *df* = 2, *P* = 0.06) (Figure 4). The mean nighttime reproductive success almost doubled between control and high ALAN conditions (3.95 ± 1.03 and 7.75 ± 1.14 offspring in average, respectively), and between low and high ALAN conditions (3.96 ± 0.82 and 7.75 ± 1.14 offspring in average, respectively).

**Figure 4.**
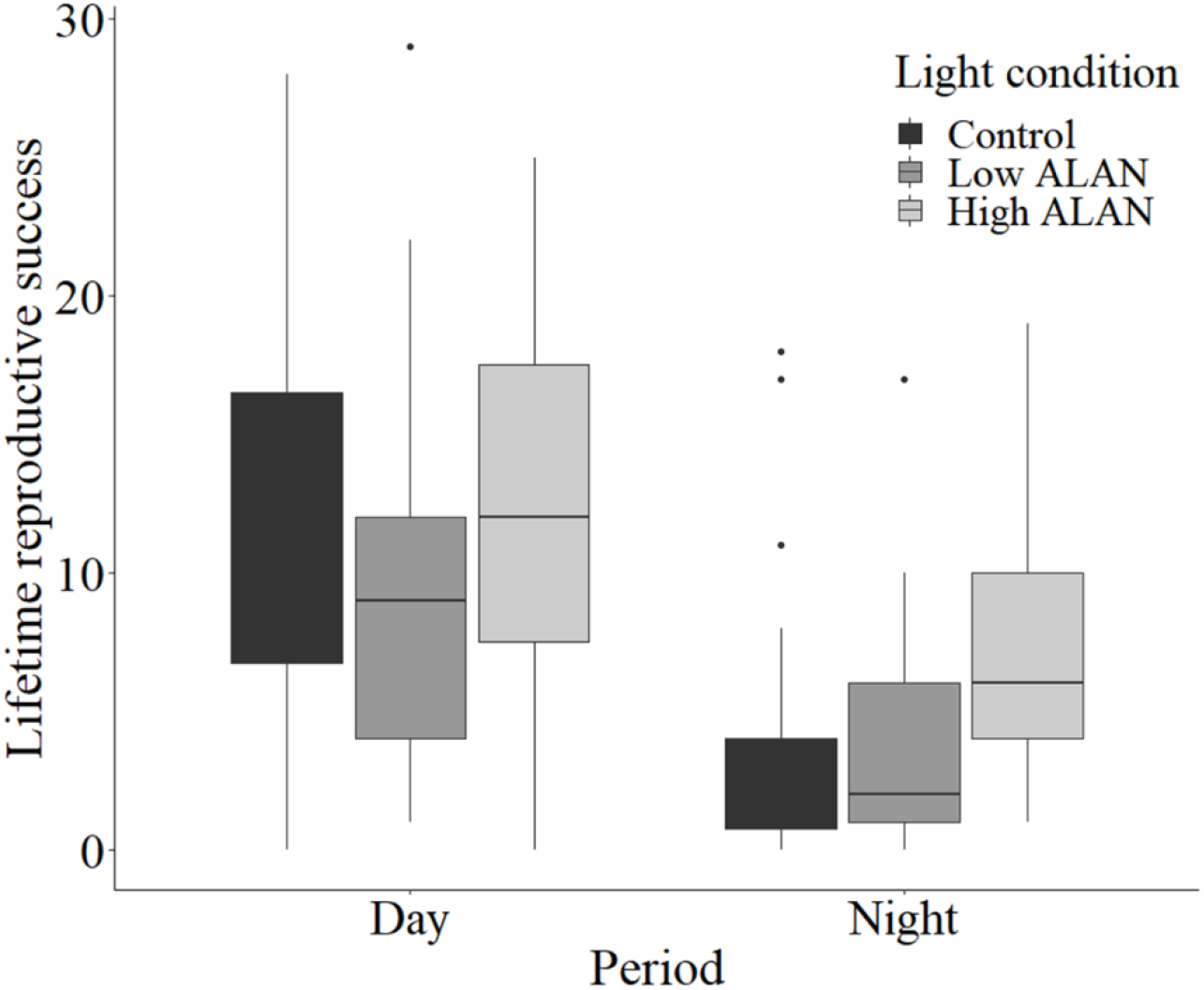
Lifetime reproductive success, estimated by the total number of offspring emerging from the host patches exploited either during the day or at night throughout the life of the wasps, for the control and artificial light at night (ALAN) conditions. The boxes represent the interquartile range (IQR) between the first quartile (Q1) and the third quartile (Q3), and the vertical lines below and above the boxes extend to the smallest (Q1 – 1.5 x IQR) and largest (Q3 + 1.5 x IQR) values, respectively. The box plot black lines represent the median lifetime reproductive success, and outliers are displayed as points.

When considering the overall lifetime reproductive success, no influence of light condition was detected (*χ* ² = 2.03, *df* = 2, *P* = 0.36). Only body size had a positive effect on the total number of offspring produced throughout a wasp’s life (*χ* ² = 4.98, *df* = 1, *P* = 0.03). However, light condition influenced the respective contribution of daytime and nighttime reproductive success to lifetime reproductive success (*F* = 3.65, *df* = 2, *P* = 0.03). Offspring coming from host patches exploited during nighttime represented 24%, 28% and 37% of the total number of offspring in the control, low ALAN and high ALAN conditions, respectively (Figure 5). Post-hoc tests showed that only the difference between high ALAN and control conditions was significant (*z* = 2.59, *P* = 0.03), but with a small effect size (Cohen’s *h* = 0.29). There was nevertheless a change in the distribution of egg-laying events between daytime and nighttime in the presence of artificial light at night.

**Figure 5.**
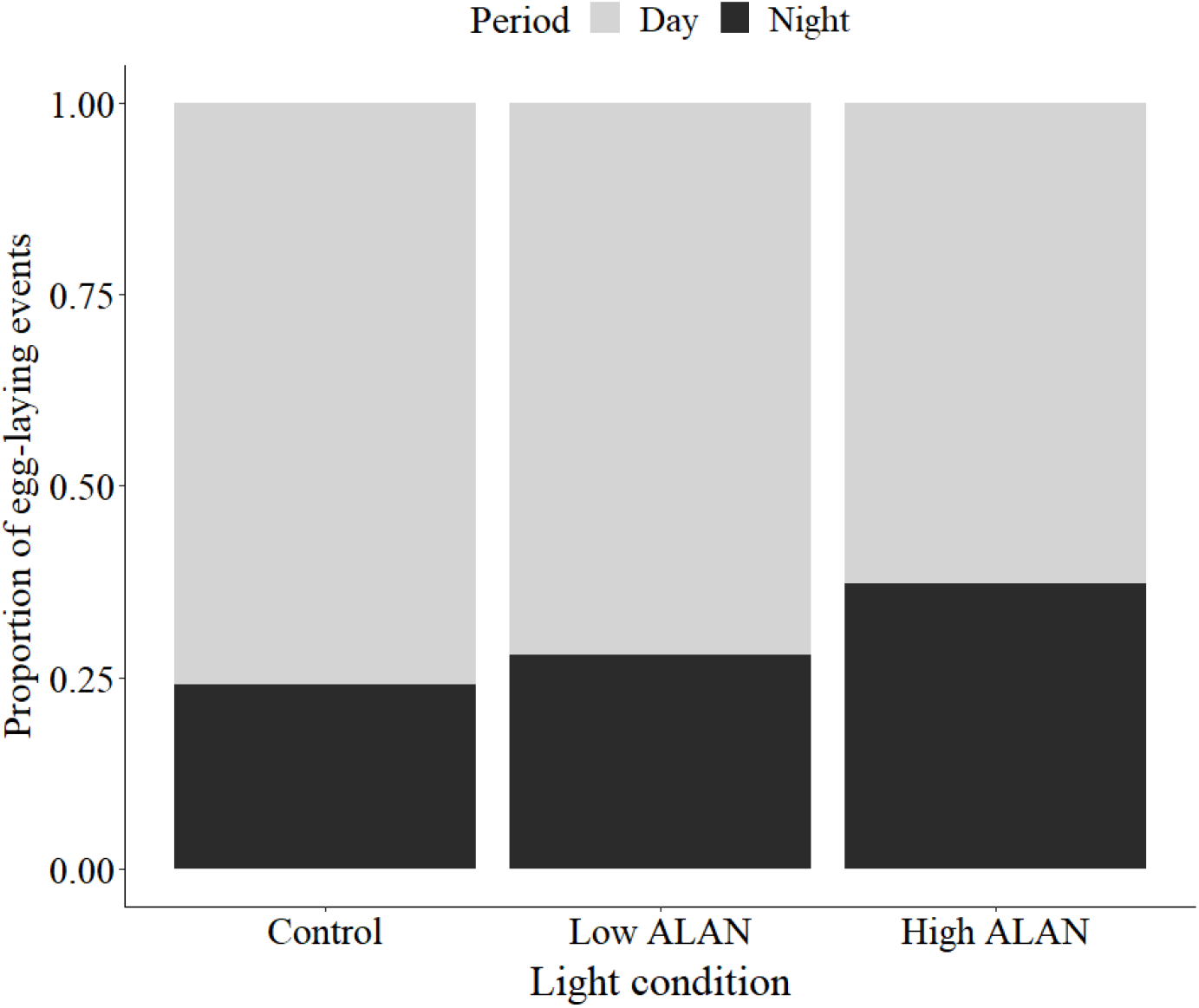
Proportion of egg-laying events that occurred during the day and during the night, illustrating the respective contribution of daytime and nighttime reproductive success to lifetime reproductive success for control and artificial light at night (ALAN) conditions.

## DISCUSSION

Our study highlights that reproductive senescence occurs in *V. canescens*, and is modulated by the degree of light exposure during the night when focusing on offspring development time. Moreover, when exposed to ALAN, the diurnal wasps were able to use the ‘night-light niche’ as illustrated by the increase in feeding and egg-laying activity at night in exposed individuals. However, the nighttime activity induced by ALAN did not affect the wasps’ lifetime reproductive success. We nevertheless detected that ALAN influenced the distribution of egg-laying events, by increasing the contribution of nighttime reproductive success to lifetime reproductive success.

As in a number of other insect species (*e*.*g*., cockroach *Nauphoeta cinerea*, Moore & Moore, 2001; or fruit flies, Carey & Molleman, 2010), we detected changes in reproductive performance with mother’s age in *Venturia canescens*. However, only age-related changes in offspring development time were impacted by artificial light at night. Specifically, offspring development time was negatively correlated with mother’s age for mothers who had been exposed to low artificial light at night. A previous study had found an interaction between mother’s age and mother’s light condition on offspring development time, but in the opposite direction (Gomes *et al*., 2021). Even though these results may at first appear contradictory, they arose from different experimental conditions. In the previous study, older females had no oviposition experience, contrary to our experiment. In *Venturia canescens*, egg maturation occurs throughout the life of the wasps (Jervis *et al*., 2001). Therefore, patch exploitation activity and presence of ALAN could have altered the quality of the eggs produced by the females. These eggs do not contain energy reserves (Le Ralec, 1995), but other substances such as hormones (*e*.*g*., melatonin whose synthesis can be influenced by ALAN (Grubisic *et al*., 2019) could be maternally transmitted to the offspring. Moreover, although a decrease in development time did not seem to impact offspring quality (estimated by the body size of the offspring at emergence) in our study, this consequence of ALAN may affect other offspring fitness-related traits (*e*.*g*., storage of energy reserves, physiological condition) that would be relevant to explore in the future. Nevertheless, evidence of relationship between light at night and reproductive senescence are rare. A study on *D. melanogaster* found that exposition to light at night influenced (non-linearly) age-related declines in the propensity to lay eggs and the number of eggs produced by females (McLay *et al*., 2018). More studies are needed to determine precisely relationships between light at night, reproductive senescence and potential underlying mechanisms.

Our experiment showed that presence of artificial light at night led *V. canescens* females to use the ‘night-light niche’. This phenomenon had rarely been reported and quantified in insect species until recently. A recent study showed that parasitism rate of an aphid parasitoid (*Aphidius megourae*) almost doubled when individuals were exposed to low levels of light at night (between 0.1 and 5 lux) (Sanders *et al*., 2018). The authors also showed that *A. megourae* did not attack hosts during the night in absence of ALAN. Another study on *A. megourae* showed that ALAN increased the parasitoid parasitism rate and intensified the positive effect of day length on the global parasitism rate (Kehoe *et al*., 2020).

However, most of the studies did not differentiate between daytime and nighttime parasitism activity in the light-polluted conditions. Even though a study showed that *V. canescens* did not move in the dark (Gomes *et al*., 2021), in our experiment wasps fed and laid eggs during the night despite the absence of light. This ‘overestimation’ of nocturnal activity may be due to the easy access to both food and hosts in our experimental set-up, whereas in field conditions, resources may be more difficult to reach in the dark, because the wasps use habitat structure to navigate (Desouhant *et al*., 2003). Nectar can be produced at night (Antoń *et al*., 2017), and visual cues are involved in foraging in *V. canescens*, with wasps being able of associative learning between colour cues and a food or host reward (Desouhant, Navel, *et al*., 2010; Lucchetta *et al*., 2008). Such visual cues probably increase the efficiency of females for finding hosts or food, but their use in absence of light might be less efficient. Moreover, temperature was held constant between day and night in our experiment, which could have exacerbated the consequences of ALAN on parasitoid activity. Indeed, while temperature has negligible effect in control conditions (Kehoe *et al*., 2020; Sanders *et al*., 2018), a drop in nighttime temperatures can moderate the effect of ALAN on parasitism rate (Kehoe *et al*., 2020). However, thelytokous *V. canescens* lives mostly in anthropogenic habitats (*e*.*g*., mills or granaries) where temperature tends to remain rather constant compared to field conditions.

These changes in activity patterns may have consequences on lifetime reproductive success. In *V. canescens*, lifetime reproductive success was not affected by artificial light at night, despite changes in feeding behaviour, intensification of nighttime egg-laying behaviour and differences in lifespan. Surprisingly, wasps exposed to high ALAN lived longer than those exposed to low ALAN in our experiment, which may be linked to the observed increase in nighttime feeding. Light at night has been shown to modify food intake in vertebrates, but also to cause various metabolic disorders (*e*.*g*., weight gain, lipid accumulation in liver, impaired glucose tolerance) (Batra *et al*., 2019; Fleury *et al*., 2020; Fonken *et al*., 2010; Masís-Vargas *et al*., 2019). Even though we did not detect a negative effect of ALAN on the lifespan of *V. canescens* in our study, changes in the timing of feeding and potential metabolism alterations due to light at night may affect fitness, and more specifically reproductive traits (Xu *et al*., 2011). Daytime reproductive success was not statistically different between light conditions, and there was only a tendency for ALAN to increase the nighttime reproductive success. This tendency was nevertheless supported by a shift in the distribution of egg-laying events between day and night for the wasps exposed to ALAN. One way to confirm or invalidate this tendency could be to provide more hosts to the wasps to allow a greater variability in the number of offspring produced and therefore detect clear differences in the reproductive success of *V. canescens* between light conditions. Several studies assessed the fitness differences under various regimes of light at night in laboratory, with various outcomes. In *Drosophila melanogaster*, a species that has an optimal oviposition behaviour at dusk or early night, chronic exposure to dim light at night reduced significantly (by 20%) the proportion of ovipositing females and the number of eggs laid (McLay *et al*., 2017). A subsequent study showed that the negative consequences of light at night persisted as females aged (McLay *et al*., 2018), confirming the probable fitness costs of ALAN in this species. However, some species seem to benefit from the presence of light at night, such as the diurnal lizard *Anolis sagrei*. In this species often found in urbanized habitats, females exposed to artificial light at night increased their reproductive output by laying more eggs at a higher rate, with no cost in terms of offspring quality (Thawley & Kolbe, 2020). Using mesocosms, a recent field experiment demonstrated that the daytime performance of the parasitoid *A. megourae* decreased markedly in the long term in presence of ALAN, suggesting that the wasps did not use the ‘night-light niche’ (even though it was not explicitly tested here) and might have impaired host finding abilities (Sanders *et al*., 2022). Interestingly, the consequences of ALAN on parasitoid fitness depended on the light wavelength. For instance, white light did not influence the parasitism rate of *A. megourae* in the laboratory (Sanders *et al*., 2022). Future studies should therefore take into account both daytime and nighttime behaviours, as well as light wavelengths, to fully apprehend the fitness consequences of ALAN on insects.

Even though ALAN is known to have multiple and pronounced detrimental consequences on individuals and populations (Boyes *et al*., 2021; Gaston *et al*., 2014), recent studies also highlighted positive effects of light at night on important biological traits at the individual level (*e*.*g*., McLay *et al*., 2018) or on species interactions. For instance, despite an overall negative effect on diurnal plant-pollinator interactions during daytime, Giavi and colleagues (2021) observed increased plant-pollinator interactions under light at night for a few plant species, which can be beneficial. In our study, although we did not detect strong detrimental nor beneficial consequences of ALAN on the lifetime reproductive success of *V. canescens*, we did observe behavioural changes when the wasps were exposed to ALAN. These changes (*i*.*e*., feeding and laying eggs at night) could result in a shift in the distribution of ovipositions between day and night for wasps which, in natural conditions, could become more vulnerable to predation or experience sub-optimal conditions for the exploitation of hosts. Behavioural changes induced by ALAN may therefore have major consequences for population dynamics and need to be explored further.

## Acknowledgements

We thank Marc Théry for lending the illuminance meter and dedicate this work to him. The research was partly funded by IDEXLyon (Scientific Breakthrough project). This work is part of the project ‘ALAN One Health’ funded by the University of Lyon 1.

