## Supplementary data for "Foraging at night under artificial light: consequences on reproductive senescence and lifetime reproductive success for a diurnal insect"

**(A)**

Bottom plate (top view)

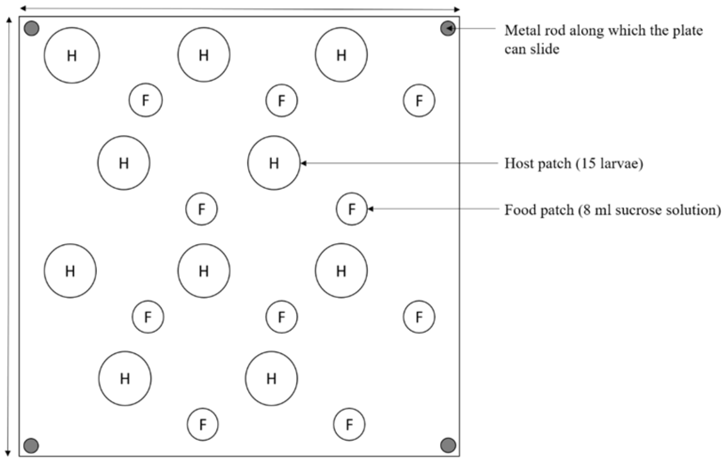


**(B)**

Superior plate (top view)

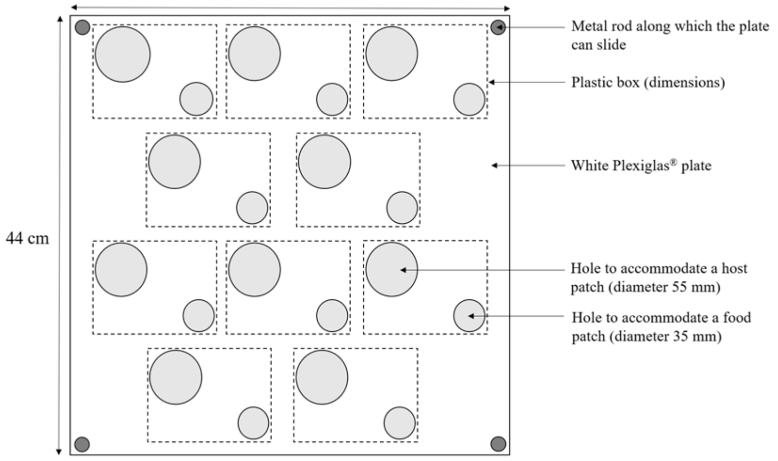


Figure S1 Diagram of the experimental set-up designed to measure the lifetime reproductive success of *V. canescens* (top view). (A) represents the bottom plate. (B) represents the superior plate, grey areas indicating empty zones (*i.e.*, holes).

**(A)** Experimental set-up (side view, lifted)

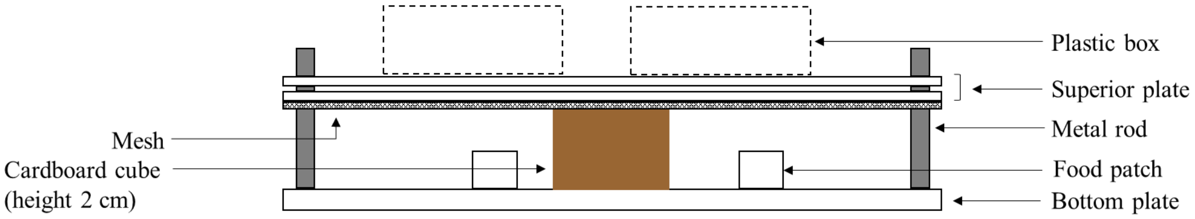


**(B)** Experimental set-up (side view, lowered)

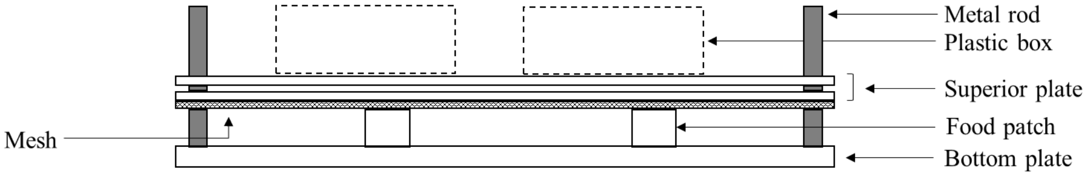


Figure S2 Diagram of the experimental set-up designed to measure the lifetime reproductive success in *Venturia canescens* (side view). (A) represents the superior plate in lifted position, which prevents the wasps to access the food and host patches. (B) represents the superior plate in lowered position, which enables the wasps to feed and parasitize host patches (not shown on the diagram).

### Description of patch replacement:

In the evening, before the end of the photophase, the superior plates were lifted to stop the wasps from accessing the patches. Host patches were replaced with new ones, and food patches were provided or removed, depending on the day (see Table 1). The mesh under the superior plate was also changed to remove any odour deposited by the wasps during the day or kairomone and food residues. Twenty minutes after the end of the photophase, the superior plates were gently lowered so that the wasps could start feeding and exploiting host patches. In the morning, access to host and food was prevented just before the onset of the photophase by lifting the superior plate again. The host patches and the mesh were replaced by new ones, and food patches were provided or removed, depending on the day. We checked if the wasps fed during the night by looking at the colour of their abdomen. Host and food were made available again for the wasps at the same time in each light condition.

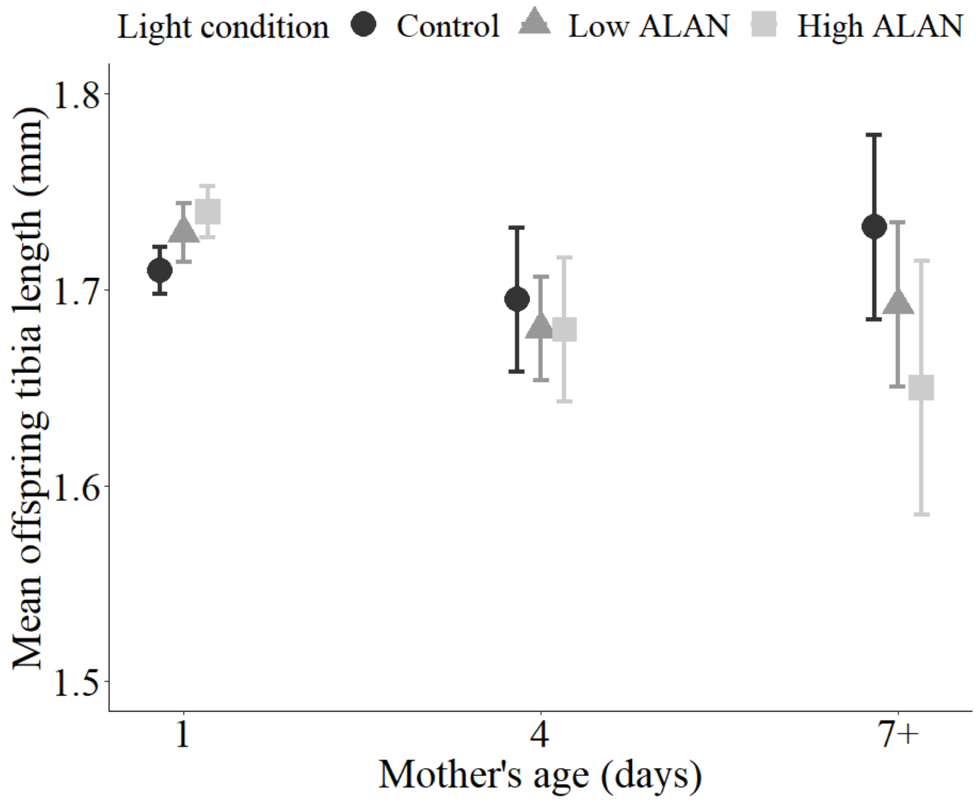


Figure S3 Effect of mother’s age on offspring body size in the control, low artificial light at night (ALAN) and high ALAN conditions. Individuals older than 7 days of age were grouped into the ‘7+’ age class. Symbols represent mean ± SE.

Table S1 Set of Cox models fitted to assess the effect of artificial light at night on lifespan in *V. canescens*. We tested the effect of light condition ('LC'), body size ('BS') and thermostated chamber ('TC'), as well as their two-by-two interactions ('LC:BS', 'LC:TC', 'BS:TC'). The symbol '+' means that the focus parameter is present in the model and 'NA' means that the focus parameter is absent from the model. 'ΔAIC' is the difference of AIC between each model and the model with the lowest AIC and 'weight' is the AIC weight of each model. The best model selected is shown in bold.

| Model ID | LC | BS | TC | LC:BS | LC:TC | BS:TC | AIC | ΔAIC | weight |
|---|---|---|---|---|---|---|---|---|---|
| 9 | + | + | + | NA | + | NA | 431.73 | 0.00 | 0.32 |
| **12** | **+** | **+** | **NA** | **NA** | **NA** | **NA** | **431.88** | **0.15** | **0.30** |
| 15 | NA | + | NA | NA | NA | NA | 434.21 | 2.48 | 0.092 |
| 11 | + | + | + | NA | NA | NA | 434.33 | 2.60 | 0.087 |
| 4 | + | + | + | NA | + | + | 435.38 | 3.65 | 0.051 |
| 3 | + | + | + | + | + | NA | 435.69 | 3.96 | 0.044 |
| 6 | + | + | NA | + | NA | NA | 435.86 | 4.13 | 0.041 |
| 7 | + | + | + | NA | NA | + | 437.25 | 5.52 | 0.020 |
| 13 | NA | + | + | NA | NA | NA | 437.42 | 5.69 | 0.019 |
| 5 | + | + | + | + | NA | NA | 438.22 | 6.49 | 0.012 |
| 1 | + | + | + | + | + | + | 439.35 | 7.62 | <0.01 |
| 8 | NA | + | + | NA | NA | + | 440.03 | 8.30 | <0.01 |
| 2 | + | + | + | + | NA | + | 441.03 | 9.30 | <0.01 |
| 10 | + | NA | + | NA | + | NA | 442.08 | 10.35 | <0.01 |
| 16 | + | NA | NA | NA | NA | NA | 444.11 | 12.37 | <0.01 |
| 18 | NA | NA | NA | NA | NA | NA | 446.40 | 14.67 | <0.01 |
| 14 | + | NA | + | NA | NA | NA | 447.33 | 15.60 | <0.01 |
| 17 | NA | NA | + | NA | NA | NA | 450.25 | 18.52 | <0.01 |

Table S2 Set of models fitted to assess the senescence pattern of daily reproductive success in *V. canescens*. We tested the effect of light condition ('LC'), 3 age functions (linear 'AL', quadratic 'AQ', threshold set at 5 days 'AT'), the interaction between light condition and age ('LC:A'), body size ('BS') and individual longevity ('LG'). The symbol '+' means that the focus parameter is present in the model and 'NA' means that the focus parameter is absent from the model. 'ΔAIC' is the difference of AIC between each model and the model with the lowest AIC and 'weight' is the AIC weight of each model. The best model selected is shown in bold.

| Model ID | LC | Age function | LC:A | BS | LG | AIC | ΔAIC | weight |
|---|---|---|---|---|---|---|---|---|
| 213 | NA | AL | NA | + | NA | 2154.63 | 0.00 | 0.50 |
| **24** | **NA** | **AL** | **NA** | **NA** | **NA** | **2156.06** | **1.43** | **0.25** |
| 182 | + | AL | NA | + | NA | 2157.19 | 2.55 | 0.14 |
| 19 | + | AL | NA | NA | NA | 2158.79 | 4.16 | 0.06 |
| 181 | NA | AL | NA | + | + | 2160.63 | 5.99 | 0.03 |
| 21 | NA | AL | NA | NA | + | 2162.28 | 7.65 | 0.01 |
| 2 | + | AL | NA | + | + | 2162.99 | 8.35 | <0.01 |
| 17 | + | AL | NA | NA | + | 2164.88 | 10.25 | <0.01 |
| 141 | + | AL | + | + | NA | 2166.62 | 11.98 | <0.01 |
| 143 | + | AL | + | NA | NA | 2168.20 | 13.57 | <0.01 |
| 14 | + | AL | + | + | + | 2172.36 | 17.73 | <0.01 |
| 142 | + | AL | + | NA | + | 2174.26 | 19.63 | <0.01 |
| 40 | NA | AQ | NA | + | NA | 2178.20 | 23.57 | <0.01 |
| 44 | NA | AQ | NA | NA | NA | 2179.22 | 24.59 | <0.01 |
| 381 | + | AQ | NA | + | NA | 2181.14 | 26.50 | <0.01 |
| 60 | NA | AT | NA | + | NA | 2182.22 | 27.59 | <0.01 |
| 39 | + | AQ | NA | NA | NA | 2182.30 | 27.67 | <0.01 |
| 38 | NA | AQ | NA | + | + | 2182.80 | 28.17 | <0.01 |
| 64 | NA | AT | NA | NA | NA | 2183.13 | 28.49 | <0.01 |
| 41 | NA | AQ | NA | NA | + | 2184.37 | 29.74 | <0.01 |
| 59 | + | AT | NA | + | NA | 2185.26 | 30.63 | <0.01 |
| 4 | + | AQ | NA | + | + | 2185.44 | 30.81 | <0.01 |
| 58 | NA | AT | NA | + | + | 2185.73 | 31.10 | <0.01 |
| 62 | + | AT | NA | NA | NA | 2186.29 | 31.66 | <0.01 |
| 37 | + | AQ | NA | NA | + | 2187.22 | 32.58 | <0.01 |
| 61 | NA | AT | NA | NA | + | 2187.25 | 32.62 | <0.01 |
| 6 | + | AT | NA | + | + | 2188.49 | 33.86 | <0.01 |
| 57 | + | AT | NA | NA | + | 2190.21 | 35.57 | <0.01 |
| 542 | + | AT | + | + | NA | 2193.47 | 38.83 | <0.01 |
| 541 | + | AT | + | NA | NA | 2194.50 | 39.86 | <0.01 |
| 54 | + | AT | + | + | + | 2196.65 | 42.02 | <0.01 |
| 543 | + | AT | + |  | + | 2198.35 | 43.72 | <0.01 |
| 342 | + | AQ | + | + | NA | 2201.28 | 46.64 | <0.01 |
| 341 | + | AQ | + | NA | NA | 2202.45 | 47.81 | <0.01 |
| 34 | + | AQ | + | + | + | 2205.52 | 50.89 | <0.01 |
| 343 | + | AQ | + | NA | + | 2207.32 | 52.68 | <0.01 |
| 211 | NA | NA | NA | + | + | 2221.45 | 66.82 | <0.01 |
| 22 | NA | NA | NA | NA | + | 2223.13 | 68.50 | <0.01 |
| 18 | + | NA | NA | + | + | 2224.17 | 69.54 | <0.01 |
| 20 | + | NA | NA | NA | + | 2226.04 | 71.41 | <0.01 |
| 25 | NA | NA | NA | + | NA | 2233.58 | 78.94 | <0.01 |
| 212 | + | NA | NA | + | NA | 2236.10 | 81.47 | <0.01 |
| 23 | + | NA | NA | NA | NA | 2237.40 | 82.76 | <0.01 |
